## supplemental figures for "Meiotic Nuclear Pore Complex Remodeling Provides Key Insights into Nuclear Basket Organization"

### Supplemental Figure 1

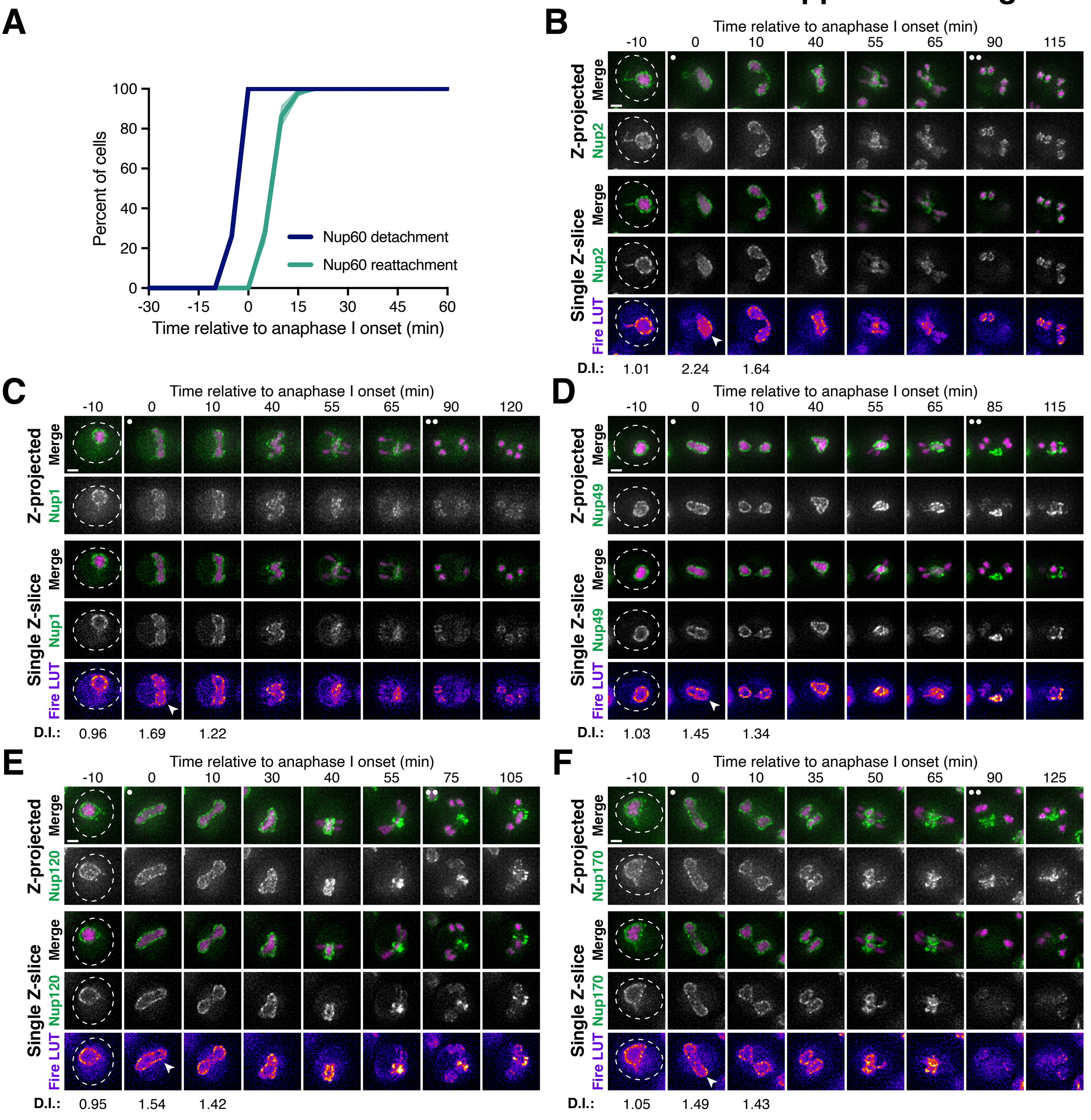

### Supplemental Figure 2

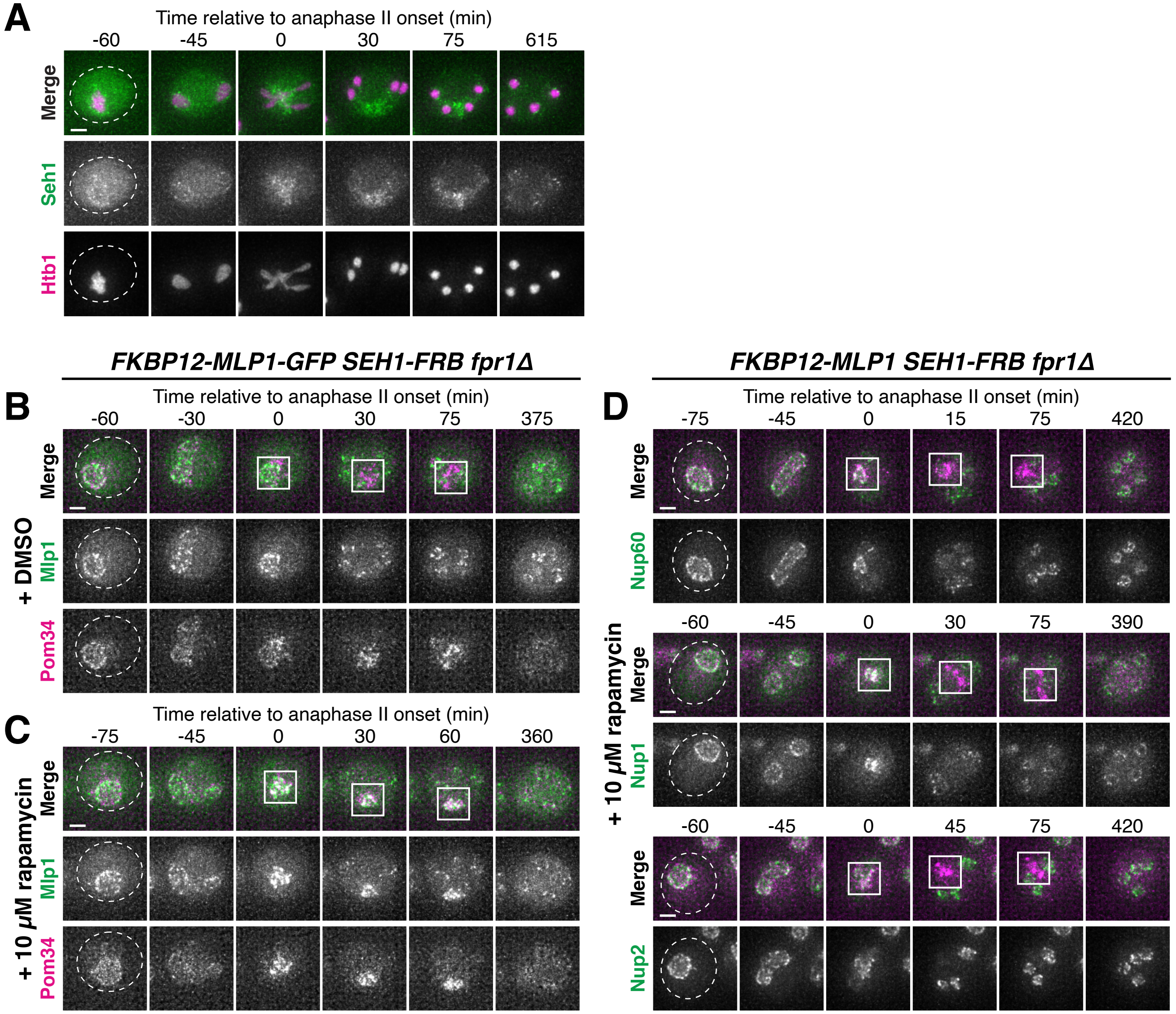

### Supplemental Figure 3

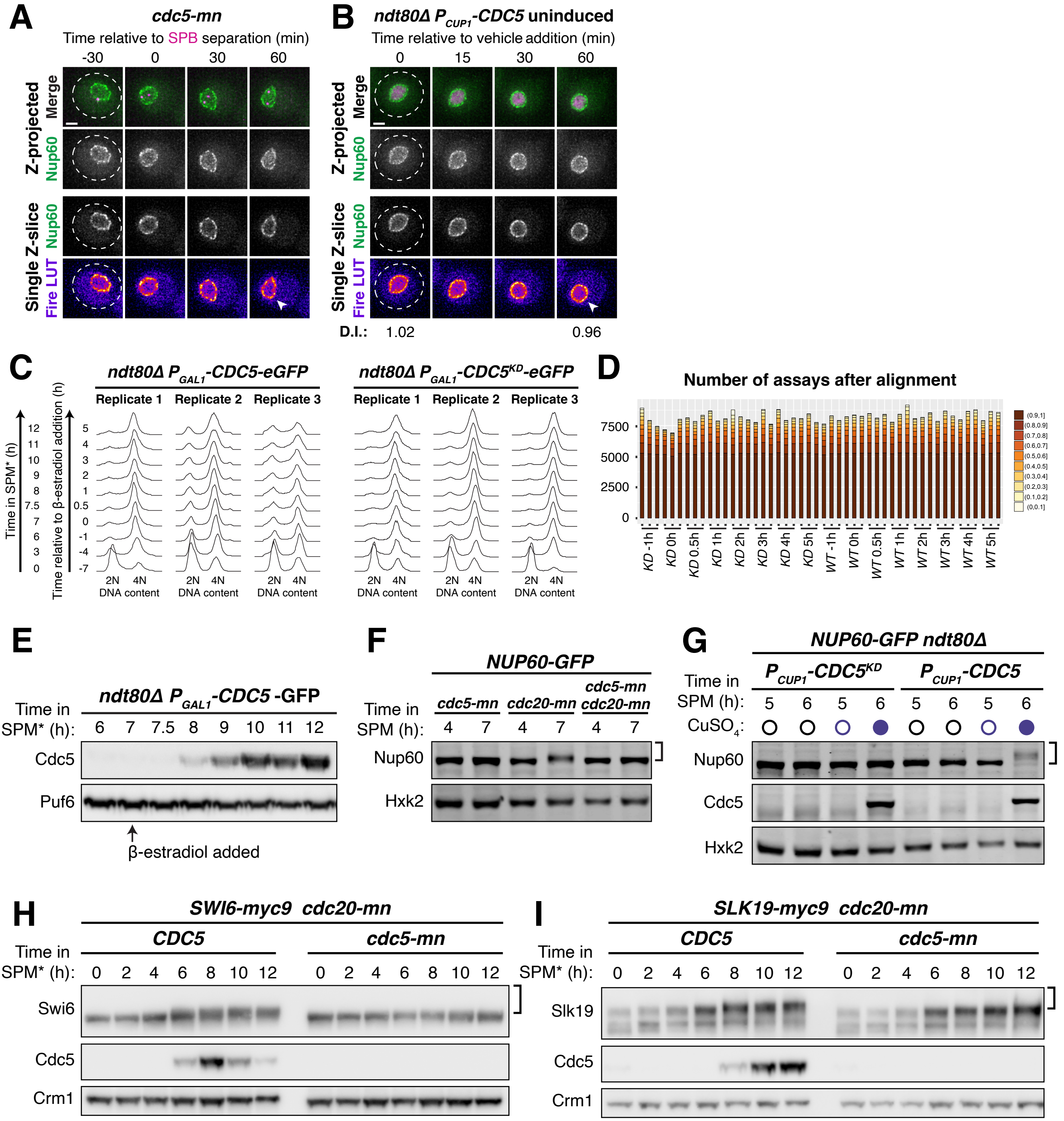

### Supplemental Figure 4

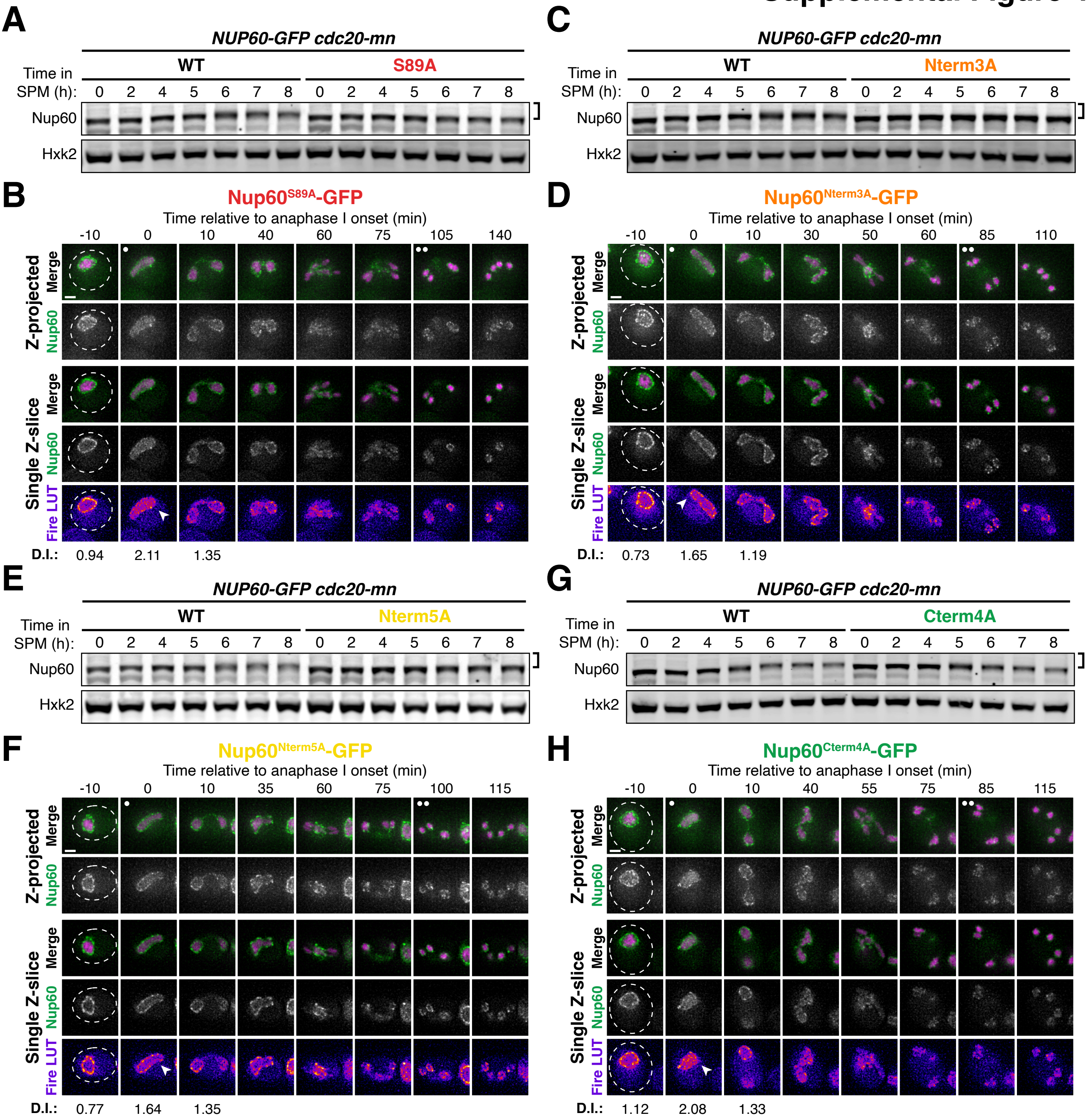

### Supplemental Figure 5

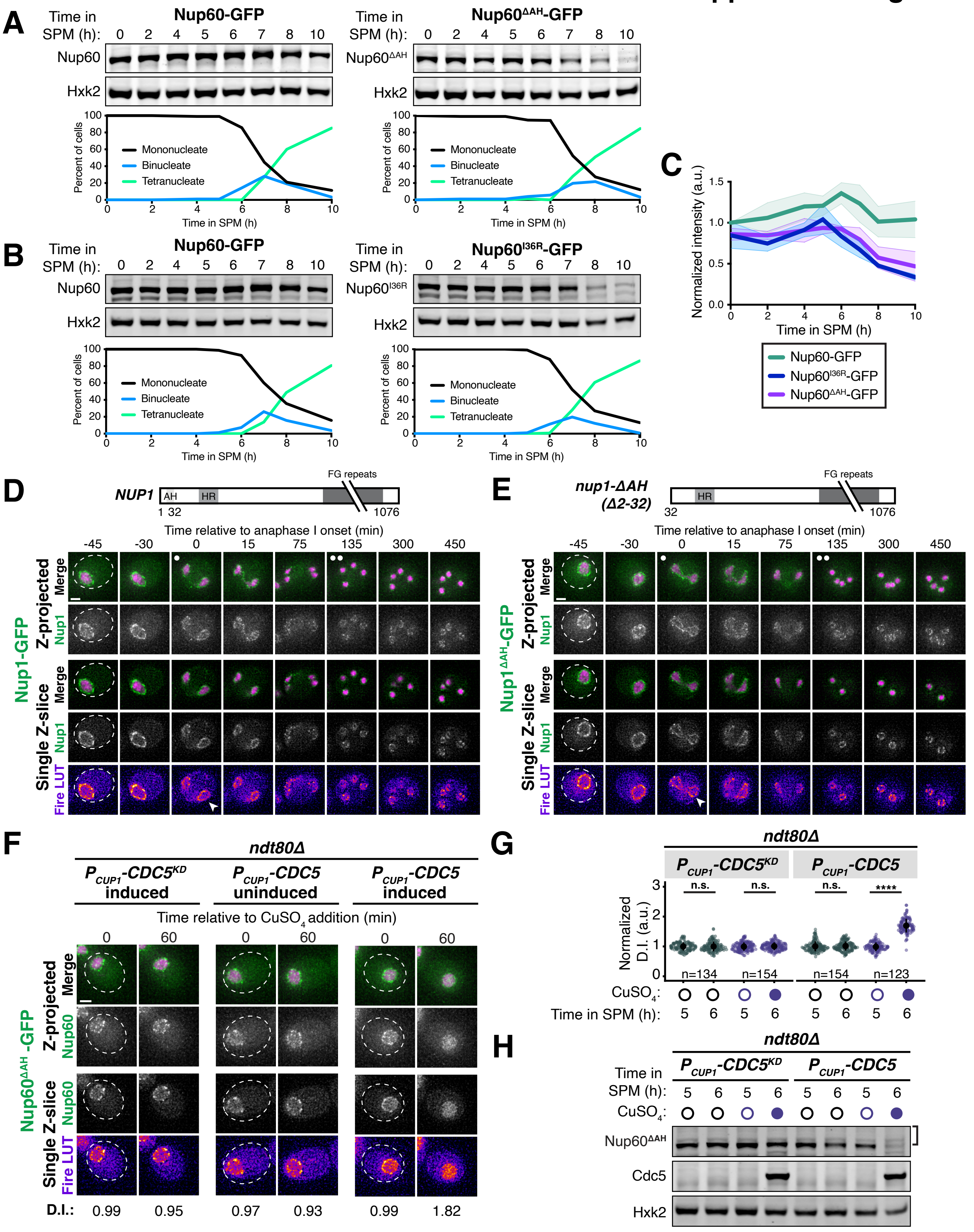

### Supplemental Figure 6

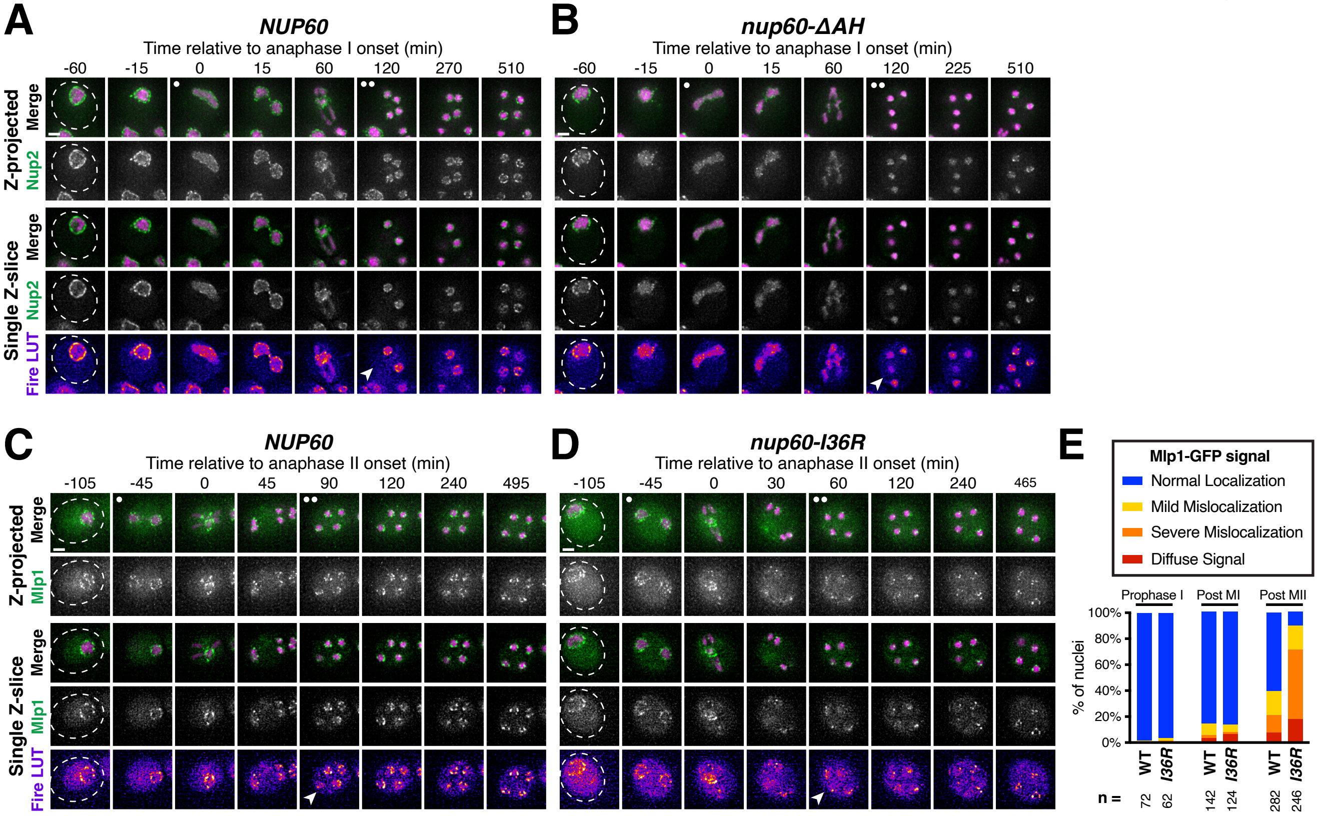

### Supplemental Figure 7

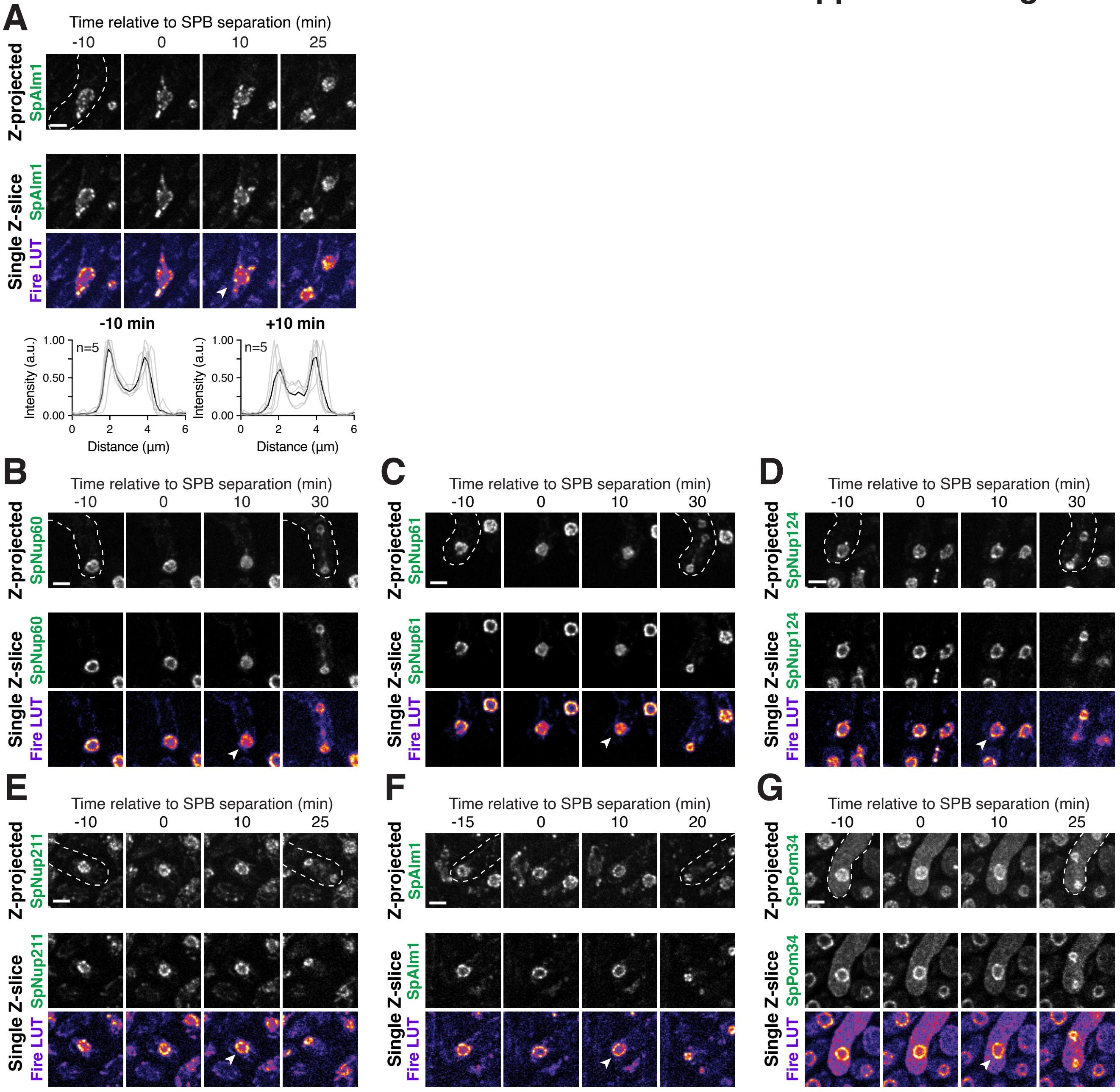
